# Tirzepatide and Intermittent Cold Exposure Independently Improve Glucose Homeostasis in DIO Mice Housed at Thermoneutrality

**DOI:** 10.64898/2026.05.21.727027

**Authors:** Annalaura Bellucci, Hadil Alfares, Clara Gale, Michael Akcan, Ben Waters, Katelyn Eisner, Bradeley Baranowski, Stewart Jeromson, Ailish Babicki-Moore, David C. Wright

**Author notes:** Corresponding author: David C. Wright, PhD, BC Children’s Hospital Research Institute, 950 West 28th Avenue, Vancouver, British Columbia, Canada, V5Z 4H4. <u>Contact for Research Governance:</u> Prof. Gail Murphy, PhD, Vice-President, Research & Innovation.

## Abstract

Obesity is considered a risk factor for metabolic diseases, including type 2 diabetes, and results from an imbalance between energy intake and energy expenditure. While pharmacological approaches such as tirzepatide, a dual GIP/GLP-1 receptor agonist, effectively reduce food intake and body weight, strategies that enhance energy expenditure (EE) may provide complementary metabolic benefits. Intermittent cold exposure (ICE) is one such approach that enhances EE and improves glucose homeostasis independent of weight loss. However, the combined effects of these interventions remain unexplored. In this study, we investigated the individual and combined effects of tirzepatide and ICE on body composition, energy metabolism, and glucose homeostasis in diet-induced obese (DIO) male and female mice housed at thermoneutrality. After 8 weeks of 45% high-fat diet feeding, mice received tirzepatide (10 nmol/kg) or vehicle and were exposed to ICE (4°C, 1 h/day, 5 days/week) or remained at thermoneutrality for 3 weeks. Energy expenditure and substrate utilization were assessed using indirect calorimetry at thermoneutrality and during an acute 1 h cold challenge. Tirzepatide reduced body weight, food intake, and adiposity in both sexes, with a greater reduction in lean mass in males. ICE did not affect body weight but improved glucose homeostasis. At thermoneutrality, tirzepatide did not alter total EE but lowered respiratory exchange ratio (RER), indicating a shift toward lipid utilization. In contrast, ICE increased energy expenditure and fat oxidation, with no additive effects observed when combined with tirzepatide. Together, these findings highlight that targeting both energy intake and expenditure represents complementary, but not necessarily additive approaches to improving metabolic health.

## Introduction

Obesity is a major risk factor for numerous chronic diseases and is associated with a range of metabolic complications, including type 2 diabetes (T2D), metabolic dysfunction-associated steatotic liver disease (MASLD), and cardiovascular disease (CVD)^1^. The pathophysiology of obesity is multifactorial, involving complex interactions between genetic, environmental, and psychological factors that contribute to overall metabolic dysregulation^2^. Over the past five decades, the global rise in obesity and its related comorbidities has intensified efforts to develop novel pharmacological interventions aimed at reducing food intake and promoting weight loss^3^. Among the current therapeutic approaches are glucagon-like peptide-1 receptor agonists (GLP-1 RAs)^4^. GLP-1 is a peptide produced by the cleavage of proglucagon and is primarily synthesized in intestinal mucosal L-cells, pancreatic islet α-cells, and neurons of the nucleus of the solitary tract.^5^ It can signal via the vagus nerve to central hypothalamic nuclei involved in regulating energy homeostasis^6^. Both orexigenic neurons (neuropeptide Y (NPY) and agouti-related peptide (AgRP) and anorexigenic neurons (proopiomelanocortin (POMC)) express the GLP-1R, allowing GLP-1 to promote satiety ^7^.

In the circulation, GLP-1 is rapidly degraded by dipeptidyl peptidase IV (DPP-IV)^8^; and therefore, degradation-resistant, long-acting GLP-1 receptor agonists have been developed for the treatment of diabetes and obesity^9^. These agents induce satiety leading to weight loss^10^ and improvements in cardiometabolic health^11^. In recent years, significant progress has been made in the development of multi-agonist therapies, including, tirzepatide, which activates both GLP-1R and the glucose-dependent insulinotropic polypeptide receptor (GIPR)^12^. Tirzepatide is considered a biased co-agonist of GLP-1R and GIPR due to its preferential stimulation of cyclic adenosine monophosphate (cAMP) production over β-arrestin recruitment, a mechanism that differs from traditional GLP-1R agonists^13^. This biased signaling results in reduced receptor internalization and increased GLP-1R expression on the cell surface, thereby enhancing its insulinotropic effects^14^. Clinical trials have demonstrated that tirzepatide leads to greater reductions in blood glucose, body weight, and triglyceride levels compared to selective GLP-1R agonists in individuals with T2D^15, 16^. Likewise, tirzepatide demonstrated superior insulin-sensitizing effects compared to the GLP-1R agonist semaglutide, independent of differences in body weight in mice with diet-induced obesity^17^.

Obesity results from an imbalance between energy intake and energy expenditure, and consequently non-pharmacological strategies to enhance energy expenditure have also been investigated. One such approach is mild cold exposure, which has shown potential in treating impaired glucose metabolism^18^. Exposure to a cold environment can increase resting energy expenditure and improve whole-body insulin sensitivity in humans^19^. In healthy individuals, the metabolic effects of cold exposure appear to be temperature-dependent, with colder stimuli producing greater reductions in blood glucose and insulin levels^18^. To further investigate the metabolic impact of cold exposure, several studies have also been conducted in rodent models, though with conflicting outcomes. Some studies reported reduced weight gain in rats exposed to cold, while others observed no significant changes in body weight^20–23^. Our group previously demonstrated that in C57BL/6J mice housed at thermoneutrality, intermittent cold exposure (ICE) improved glucose tolerance without reducing body weight^24^ suggesting an uncoupling between weight loss and improvements in glucose metabolism. Based on these findings, combining an agent that reduces food intake with an intervention that enhances energy expenditure and glucose metabolism may produce greater beneficial effects on metabolic dysfunction than either approach alone. Therefore, the aim of this study was to investigate the individual and combined effects of tirzepatide and intermittent cold exposure on weight gain, adiposity, and glucose metabolism in DIO mice housed at thermoneutrality.

## Methods

### Animal and ethics

All protocols were approved by the University of British Columbia Animal Care Committee (protocol #A22-0011) and conducted in accordance with Canadian Council on Animal Care Guidelines. Male and female C57BL/6J mice (12 weeks of age) (CAT# 000664, Jackson Laboratories) were group-housed (∼4/cage) at room temperature (20–22°C) with ad libitum access to standard chow (CAT# 2918, Teklad) and water while acclimating to the animal facility at BC Children’s Hospital Research Institute for 1 week. Following acclimation, mice were group-housed (∼4/cage) in shoe-box cages, provided ad libitum access to a 45% high-fat diet (HFD) (45% kcal fat, 35% kcal carbohydrate, 20% kcal protein) (CAT# D12451, Research Diets Inc.) and water, and maintained at thermoneutrality (29°C) in a Solace Zone Caging System (Alternative Design Manufacturing & Supply Inc.).

### Experimental outline

After 8 weeks of feeding a 45% HFD to induce obesity, male and female mice were weight-matched and assigned to the following groups: TRZ TN (tirzepatide thermoneutrality), VEH TN (vehicle thermoneutrality), TRZ ICE (tirzepatide intermittent cold exposure), and VEH ICE (vehicle intermittent cold exposure) (Figure 1A). Mice in the ICE groups underwent intermittent cold exposure (1 h/day, 5 days/week, ∼9 AM) for 3 weeks. ICE consisted of removing mice from their home cages and transferring them to pre-chilled cages, singly housed, inside a refrigerator maintained at 4°C (ESBE scientific industries Inc.). Mice in the thermoneutrality groups remained in their home cages. Following cold exposure, mice were returned to thermoneutrality. Tirzepatide (10 nmol/kg BW)^25, 26^ (CAT# 39748, Cayman Chemicals) or saline was administered daily via subcutaneous injection with an insulin syringe (5 days/week for 3 weeks), approximately 3 hours before the onset of the dark cycle (∼6 PM). This dose has previously been shown to cause weight loss and improve glucose homeostasis in DIO mice^25, 26^.

**Figure 1:**
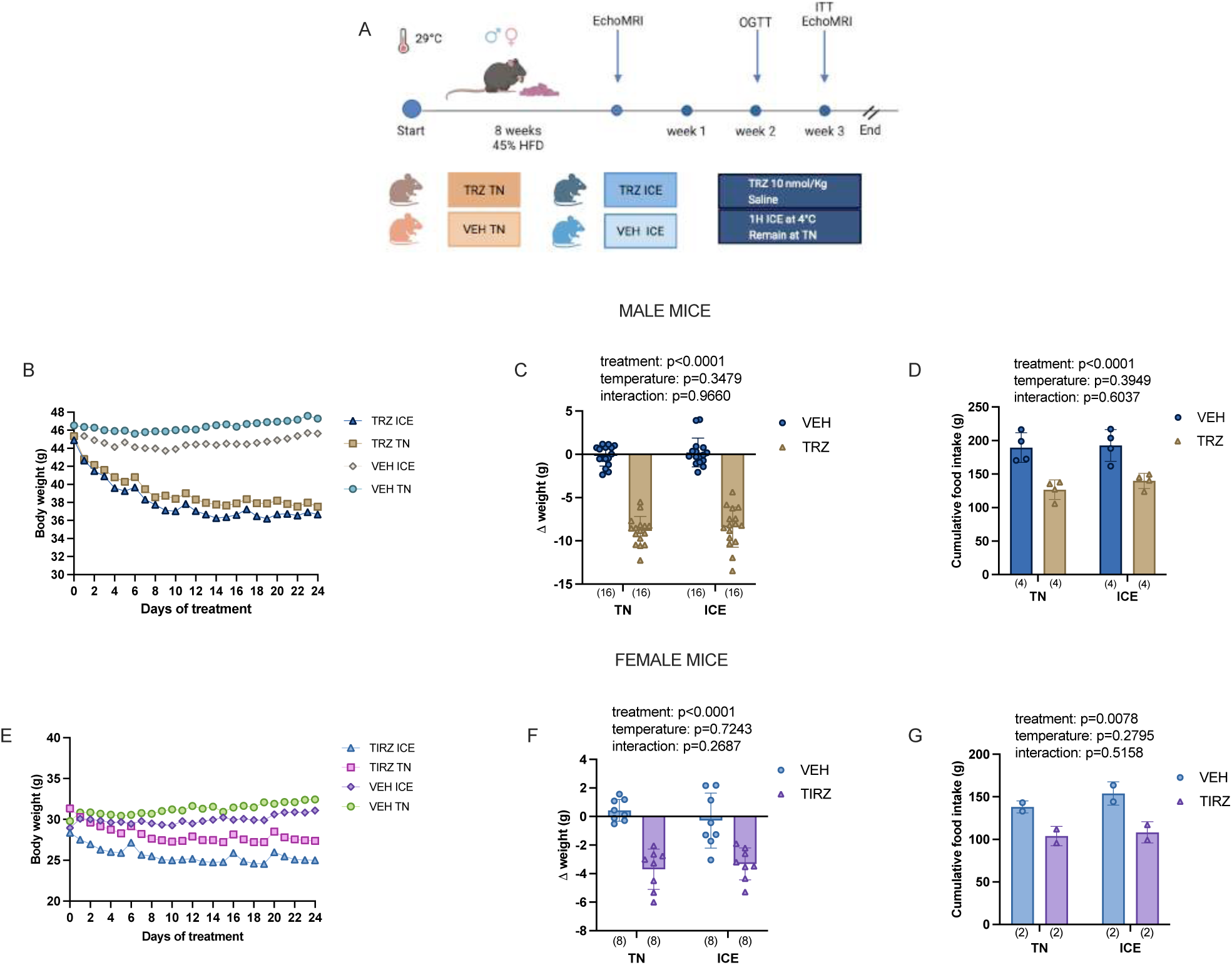
Tirzepatide, but not ICE, reduces body weight and food intake in male and female DIO mice housed at thermoneutrality. Experimental design and timeline (A). Tirzepatide reduced body weight in male (B, C) and female (E, F) mice in parallel with reductions in food intake (D,G). Statistical analysis was performed using two-way ANOVA (treatment x temperature), with main and interaction effects shown in the inset. Data are presented as mean ± SD, with individual data points shown. For clarity, mean values only are shown in B and E. Sample sizes per group are indicated in parentheses below each bar.

### Body composition and tissue collection

Pre-and post-intervention, body composition was measured using an EchoMRI body composition analyzer (EchoMRI, Houston, TX). Tissue collection occurred ∼24 h after the final cold exposure bout and tirzepatide injection, following a 4-h fast. Mice were anesthetized with pentobarbital (Euthansol; 120 mg/kg BW, i.p.) and euthanized via exsanguination. Blood was collected by cardiac puncture, kept on ice for ∼30 min, followed by centrifugation (1500 x g for 10 min) to obtain serum. The liver, inguinal white adipose tissue (iWAT), epididymal white adipose tissue (eWAT), gonadal white adipose tissue (gWAT), interscapular brown adipose tissue (iBAT), triceps, tibialis anterior, and gastrocnemius muscles were harvested, snap-frozen in liquid nitrogen, and stored at −80°C until further analyses.

### Oral glucose and insulin tolerance tests

Glucose and insulin tolerance were assessed 24 h after the last cold exposure bout and tirzepatide injection. The oral glucose tolerance test (OGTT) was performed during the second week of the intervention. Mice were fasted for 6 h (starting at ∼07:00 AM) and administered an oral gavage of glucose (2 g/kg body weight)^27^. Blood glucose was measured from a small drop collected from the tail vein using a handheld glucometer and glucose strips (Freestyle Lite; Abbott Laboratories, Abbott Park, IL, USA) immediately before gavage (time 0) and at 15, 30, 60, 90, and 120 min post-gavage. The insulin tolerance test (ITT) was performed during the third week of the intervention, one week after the OGTT^28^. Mice were fasted for 4 h (starting at ∼09:00 AM) and received an intraperitoneal bolus of human insulin (0.50 U/kg body weight; Eli Lilly, Indianapolis, IN, USA). Blood glucose was measured immediately before injection (time 0) and at 15, 30, 60, 90, and 120 min post-injection.

### Serum and tissue analyses

Serum, tissue metabolites and hormones were quantified using a Versamax Tunable Microplate Reader and SoftMax Pro Software (Molecular Devices). Insulin was measured using a commercially available enzyme-linked immunosorbent assay (ELISA) kit (CAT# 10-1247-01, Mercodia Inc.). Serum non-esterified fatty acids (NEFA) (CAT# CA97000-012, Wako Chemicals), beta-hydroxybutyrate (BHB) (CAT# 700190, Cayman Chemicals), cholesterol (CAT# 9003100, Cayman Chemicals), and triglycerides (CAT# 10010303, Cayman Chemicals) were assessed using commercially available colorimetric assay kits. Liver triglycerides were quantified using a commercially available colorimetric triglyceride assay kit (CAT# 10010303, Cayman Chemicals) following the manufacturer’s instructions.

### Metabolic cages

An additional cohort of 12-week-old C57BL/6J male mice were fed a 45% high-fat diet (HFD) to induce obesity and were housed at thermoneutrality (29 °C). After 8 weeks, mice received daily injections of tirzepatide (10 nmol/kg) or saline 5 days per week for 2 weeks and underwent daily ICE sessions at 4 °C or remained at thermoneutrality (29 °C) (Figure 6A). Mice were individually housed in metabolic cages (Promethion High-Definition Multiplexed Respirometry System for Mice; Sable Systems International, Las Vegas, NV, USA) maintained at thermoneutrality (29 °C), with ad libitum access to food and water. Mice were allowed to acclimate for 24 h before data collection over a full light–dark cycle. During the metabolic cage measurements, mice continued to receive their respective daily injections of tirzepatide or saline. Following 48 h, mice in the ICE group underwent a 1 h cold challenge (4 °C) in the metabolic cages and were euthanized immediately afterward. Mice maintained at thermoneutrality were euthanized at the corresponding time point. Respirometry measurements were recorded every 5 min, with a 30 s dwell time per cage and a baseline cage sampling frequency of 30 s every four cages. The respiratory exchange ratio (RER) was calculated as the ratio of VCO₂ to VO₂. Locomotor activity was determined based on beam breaks detected by a grid of infrared sensors in each cage. Energy expenditure was calculated using the Weir equation^29^: Energy expenditure (kcal) = 3.941 × VO₂ (L) + 1.106 × VCO₂ (L). VO₂ and VCO₂ were used to calculate fat and carbohydrate oxidation during both the 24h thermoneutrality period and the acute 1h thermoneutrality/cold exposures.

For the 24h thermoneutrality analysis, VO₂ and VCO₂ were averaged over the final 24 h at thermoneutrality, values were converted to L·min⁻¹ and fat oxidation and carbohydrate oxidation were calculated as: fat oxidation (mg·h⁻¹) = 1.695x(VO₂, L·min⁻¹ x 60) − 1.701x(VCO₂, L·min⁻¹ x 60), carbohydrate oxidation (mg·h⁻¹) = 4.585×(VCO₂, L·min⁻¹ x 60) − 3.226x(VO₂, L·min⁻¹ x 60).^30^. For the acute 1h exposures, VO₂ and VCO₂ were recorded at 1-min resolution (mL·min⁻¹), converted to L·min⁻¹. Fat oxidation and carbohydrate oxidation were calculated as: fat oxidation (mg·h⁻¹) = 1.695x(VO₂x 60) − 1.701x(VCO₂x 60)], carbonhydrate oxidation = 4.585x(VCO₂, L·min⁻¹ x 60) − 3.226x(VO₂, L·min⁻¹ x 60).

### Histology

Liver, iWAT, eWAT, and gWAT tissues were collected and fixed overnight in 10% formalin (catalog no. HT501128; Sigma, St. Louis, MO, USA). Histological processing was performed by the core histology service at BC Children’s Hospital Research Institute (Vancouver, BC, CA). Briefly, samples were dehydrated, paraffin-embedded, and sectioned at 5 μm thickness. Sections were stained with hematoxylin and eosin (HCE) and imaged on a BX61 microscope (Olympus, Tokyo, Japan) at 20x magnification. Adipocyte morphology was assessed in Fiji (ImageJ; version 2.0.2) using standardized fields of view (750×450 µm) selected from representative iWAT, eWAT and gWAT regions. Approximately 200 adipocytes per field were manually outlined using the freehand selection tool, and cross-sectional area and Feret’s diameter were recorded.

### Statistical analysis

Statistical analyses were performed using GraphPad Prism v.10.0 (GraphPad Software, La Jolla, CA, USA). Data were analyzed by two-way ANOVA, and Tukey’s post-hoc test was applied when a significant interaction between tirzepatide and temperature was detected. Associations between body weight and lean mass and body weight and fat mass were examined using Pearson or Spearman correlation coefficients, as appropriate. Data from 24h metabolic cages at thermoneutrality were processed with CalR^31^, and energy expenditure (EE) was evaluated using body weight as a covariate. Energy expenditure, fat oxidation, and carbohydrate oxidation during the 1h cold challenge were analyzed in R (version 4.5.2) using a linear regression model that included treatment, temperature, their interaction, and body weight as a covariate (EE ∼ Treatment x Temperature + Weight; Fat oxidation_mg_h_mean ∼ Treatment x Temperature + Weight; Carb oxidation ∼ Treatment x Temperature + Weight). Estimated marginal means and planned pairwise comparisons were obtained using the emmeans package with Bonferroni adjustment. Data are presented as mean ± SD, with individual data points shown when possible. Statistical significance was set at p < 0.05.

## Results

### Tirzepatide, but not ICE, reduces body weight and food intake in male and female DIO mice housed at thermoneutrality

We aimed to determine the individual and combined effects of tirzepatide and ICE on body weight and food intake in DIO male and female mice housed at thermoneutrality. Tirzepatide, but not ICE, resulted in reductions in body weight in male (figure 1B and 1C) (treatment: p < 0.0001; temperature: p = 0.3479; interaction: p = 0.9660) and female (figure 1E and 1F) (treatment: p < 0.0001; temperature: p = 0.7243; interaction: p = 0.2687) mice. Furthermore, cumulative food intake was also decreased with tirzepatide in male (figure 1D) (treatment: p < 0.0001; temperature: p = 0.3949; interaction: p = 0.6073) and female (figure 1G) (treatment: p = 0.0078; temperature: p = 0.2795; interaction: p = 0.5158) mice.

### Tirzepatide reduces fat mass and has sexually dimorphic effcts on lean mass

Incretin agonists, including tirzepatide, have been shown to reduce both lean and fat mass^32,33^. In the current, tirzepatide reduced lean mass in male mice (figure 2A) (treatment: p < 0.0001; temperature: p = 0.6025; interaction: p = 0.0920). Delta lean mass was also reduced with tirzepatide and ICE (figure 2B) (treatment: p < 0.0001; temperature: p = 0.0014; interaction: p = 0.0569) in male mice. In female mice, there was a significant treatment x temperature interaction (figure 2L) (treatment: p = 0.0661; temperature: p = 0.5662; interaction: p = 0.0085), with TRZ ICE mice exhibiting lower lean mass compared to VEH ICE (p = 0.0116) (figure 2L). Delta lean mass displayed a significant interaction (treatment: p = 0.2406; temperature: p = 0.3178; interaction: p = 0.0432), although post-hoc comparisons were not different between groups (figure 2M). Pearson correlation analysis revealed a positive correlation between lean mass and body weight in both male (r = 0.5287; p < 0.0001) (figure 2C) and female (r = 0.3588; p = 0.0437) mice (figure 2N). In male mice, tibialis anterior weight was reduced with tirzepatide (figure 2G) (treatment: p < 0.006; temperature: p = 0.0521; interaction: p = 0.8551), with a trend toward reduced gastrocnemius muscle mass (figure 2H) (treatment: p = 0.0518; temperature: p = 0.9548; interaction: p = 0.2098). No differences were observed in triceps muscle weight (figure 2I) (treatment: p = 0.1151; temperature: p = 0.8451; interaction: p = 0.5894).

**Figure 2:**
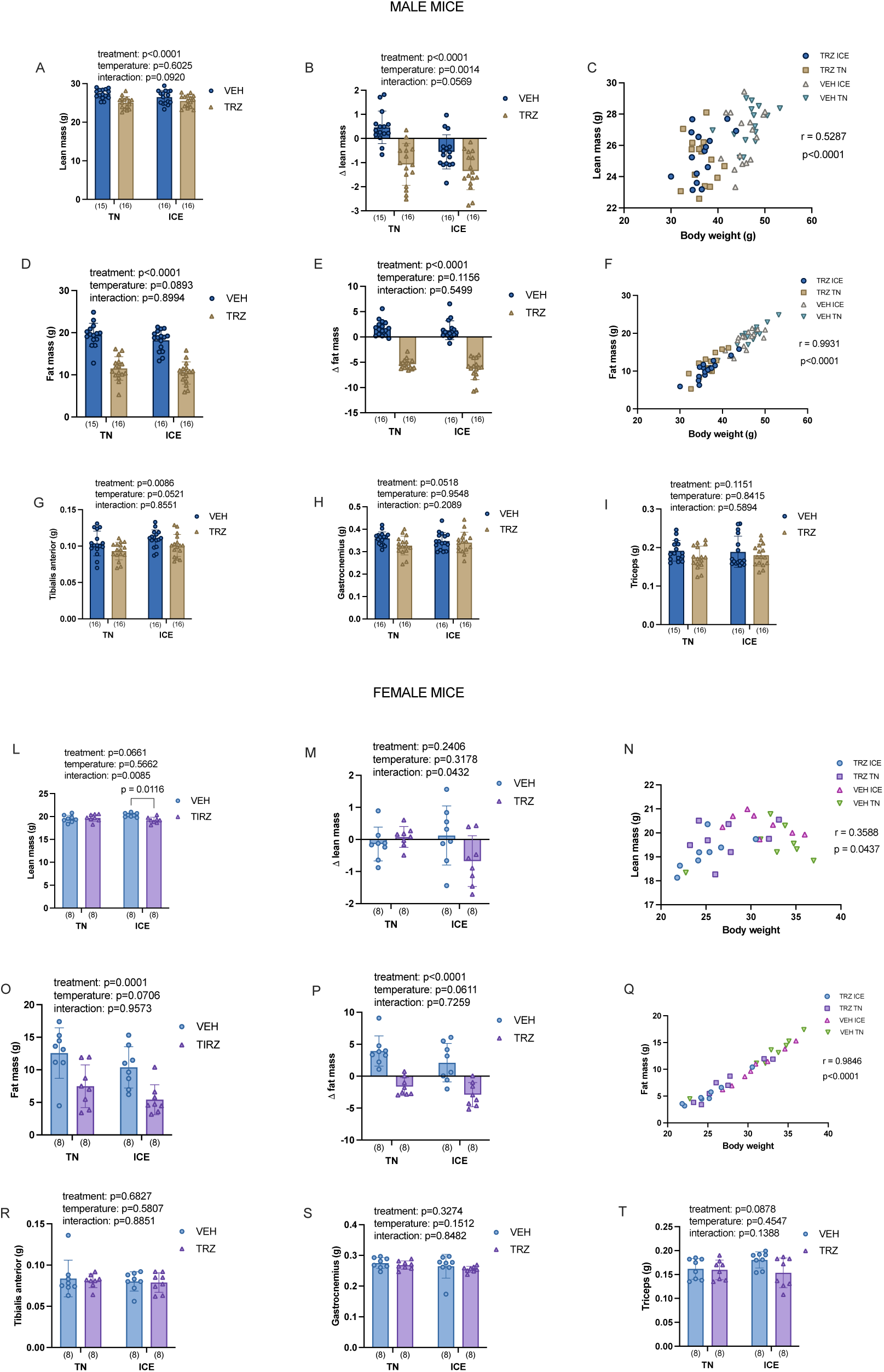
Tirzepatide reduces fat mass and has sexually dimorphic effects on lean mass. Tirzepatide reduces lean mass in male mice (A, B) and in female mice exposed to ICE (L, M). Fat mass is reduced by tirzepatide in male (D, E) and female mice (O, P). Lean (C,N) and fat mass (F, Q) positively correlates with body weight in male and female mice. Tibialis anterior (G), gastrocnemius (H), but not triceps (I) weight is reduced with tirzepetide treatment in male mice. Statistical analysis was performed using two-way ANOVA (treatment x temperature), with main and interaction effects shown in the inset. Correlation analyses were performed using Pearson or Spearman coefficients. Data are presented as mean ± SD, with individual data points shown. Sample sizes per group are indicated in parentheses below each bar.

In female mice, no differences were observed in tibialis anterior (figure 2R) (treatment: p = 0.6827; temperature: p = 0.5807; interaction: p = 0.8551), gastrocnemius (figure 2S) (treatment: p = 0.3274; temperature: p = 0.1512; interaction: p = 0.8482), or triceps muscles weight (figure 2T) (treatment: p = 0.0878; temperature: p = 0.4547; interaction: p = 0.1388). Tirzepatide reduced fat mass in both male (figure 2D) (treatment: p < 0.0001; temperature: p = 0.0893; interaction: p = 0.8994) and female (figure 2O) (treatment: p < 0.0001; temperature: p = 0.0706; interaction: p = 0.9573) mice. Delta fat mass was also significantly reduced with tirzepatide in male (figure 2E) (treatment: p < 0.0001; temperature: p = 0.1156; interaction: p = 0.5499) and female animals (figure 2P) (treatment: p < 0.0001; temperature: p = 0.0611; interaction: p = 0.7259). Spearman and Pearson correlation analyses showed a positive relationship between fat mass and body weight in male (figure 2F) (r = 0.9931; p < 0.0001) and female (figure 2Q) (r = 0.9846; p < 0.0001) mice. Overall, Tirzepatide reduced lean mass in a sexually dimorphic manner, with male mice displaying greater reduction in lean mass, whereas fat mass was robustly reduced in both sexes.

### Tirzepatide and ICE differentially affect white and brown adipose tissue mass

We found that tirzepatide decreased iWAT (figure 3A) (treatment: p < 0.0001; temperature: p = 0.1524; interaction: p = 0.3971) and eWAT mass (figure 3B) (treatment: p < 0.0001; temperature: p = 0.1180; interaction: p = 0.1604) in male mice. In female mice, both tirzepatide and ICE reduced iWAT (figure 3D) (treatment: p < 0.0001; temperature: p = 0.0144; interaction: p = 0.2921) and gWAT mass (figure 3E) (treatment: p = 0.0002; temperature: p = 0.0188; interaction: p = 0.2710). In male mice, we observed a treatment x temperature interaction in iBAT mass (figure 3C) (treatment: p < 0.0001; temperature: p < 0.0001; interaction: p = 0.0122). Specifically, VEH ICE displayed greater iBAT mass compared to all other groups (p<0.0001). iBAT mass was less in TRZ TN compared to VEH TN (p=0.0414) and TRZ ICE (p=0.0350) group. In female mice, we found a treatment x temperature interaction in iBAT mass (figure 3F) (treatment: p = 0.1072; temperature: p < 0.0001; interaction: p = 0.0147). Specifically, VEH ICE displayed higher BAT mass compared to TRZ ICE (p = 0.0262), with VEH ICE mice also showing higher iBAT mass compared to TRZ TN (p = 0.0004) and VEH TN (p < 0.0001).

**Figure 3:**
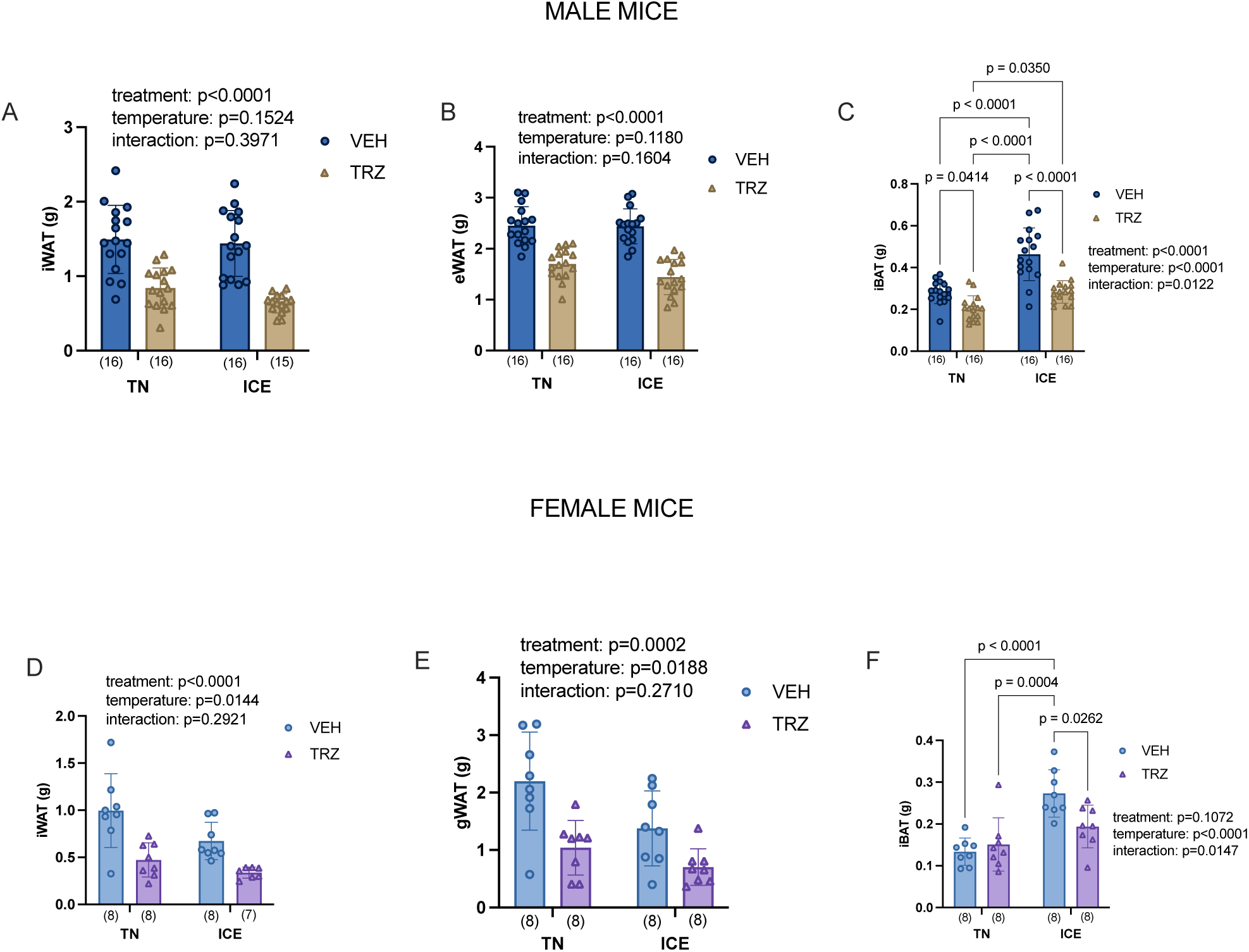
Tirzepatide and ICE differentially affect white and brown adipose tissue mass. Tirzepatide reduces iWAT mass (A) and eWAT mass (B) in male mice. In female mice, tirzepatide and ICE reduce iWAT mass (D) and and gWAT mass (E). ICE increases iBAT mass in male (C) and female mice (F). Statistical analysis was performed using two-way ANOVA (treatment x temperature), with main and interaction effects shown in the inset. Data are presented as mean ± SD, with individual data points shown. Sample sizes per group are indicated in parentheses below each bar.

### Tirzepatide affects adipocyte morphology

H&E staining revealed that adipocyte size was reduced with tirzepatide in male iWAT (figure 4A, figure 4C) (treatment: p = 0.0004; temperature: p = 0.7666; interaction: p = 0.3904) and eWAT (figure 4D, figure 4B) (treatment: p = 0.0005; temperature: p = 0.5515; interaction: p = 0.1420). In female mice, HCE staining showed that tirzepatide reduced adipocyte areas in iWAT (figure 4E, figure 4G) (treatment: p = 0.0488; temperature: p = 0.8516; interaction: p = 0.4497) and gWAT (figure 4F, figure 4H) (treatment: p = 0.0419; temperature: p = 0.1177; interaction: p = 0.7633).

**Figure 4:**
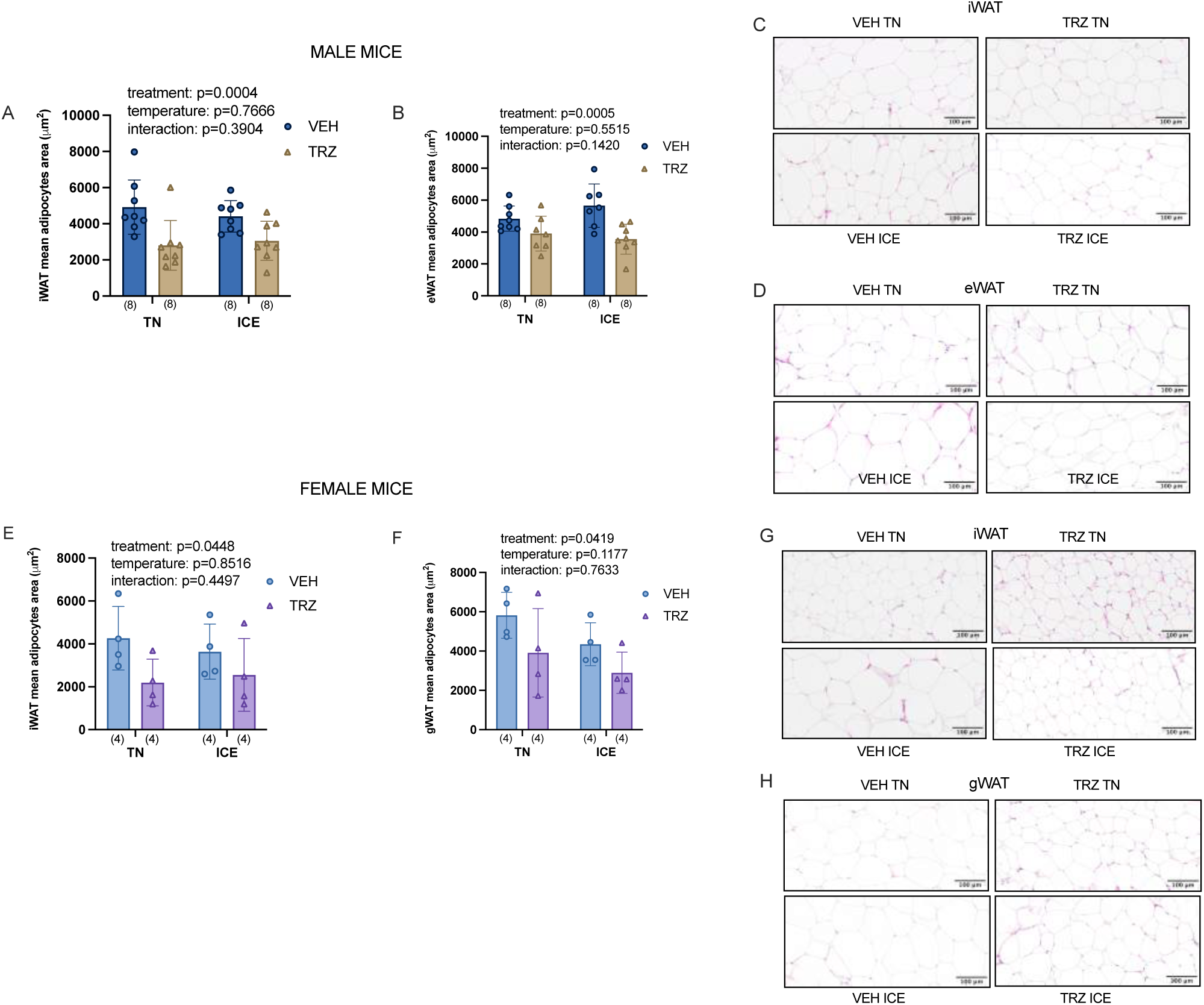
Tirzepatide affects adipocyte morphology. Tirzepatide reduces iWAT (A) and eWAT mean adipocytes area (B) in male mice. In female mice, tirzepatide reduces iWAT mean adipocytes area (E), while tirzepatide reduces gWAT mean adipocytes area (F). Representative HCE staining images are shown for iWAT (C) and eWAT (FD in male mice, and for iWAT (G) and gWAT (H) in female mice. Statistical analysis was performed using two-way ANOVA (treatment x temperature), with main and interaction effects shown in the inset. Data are presented as mean ± SD, with individual data points shown. Sample sizes per group are indicated in parentheses below each bar.

### Tirzepatide reduces liver mass and triglycerides in male mice

Treatment with tirzepatide has been shown to reduce liver fat accumulation^34^. We found that in male mice tirzepatide reduced liver mass (figure 5A) (treatment: p < 0.0001; temperature: p = 0.1963; interaction: p = 0.1238) and liver TAG (figure 5B) (treatment: p < 0.0001; temperature: p = 0.4178; interaction: p = 0.3242). In female mice, tirzepatide lowered liver mass (figure 5D) (treatment: p < 0.0001; temperature: p = 0.1065; interaction: p = 0.2512), while liver TAG levels remained unchanged (figure 5E) (treatment: p = 0.7539; temperature: p = 0.1261; interaction: p = 0.8148). Representative HCE-stained liver sections revealed qualitatively greater lipid deposition in VEH compared to tirzepatide TRZ-treated mice of both sexes (Figure 5C, 5F).

**Figure 5:**
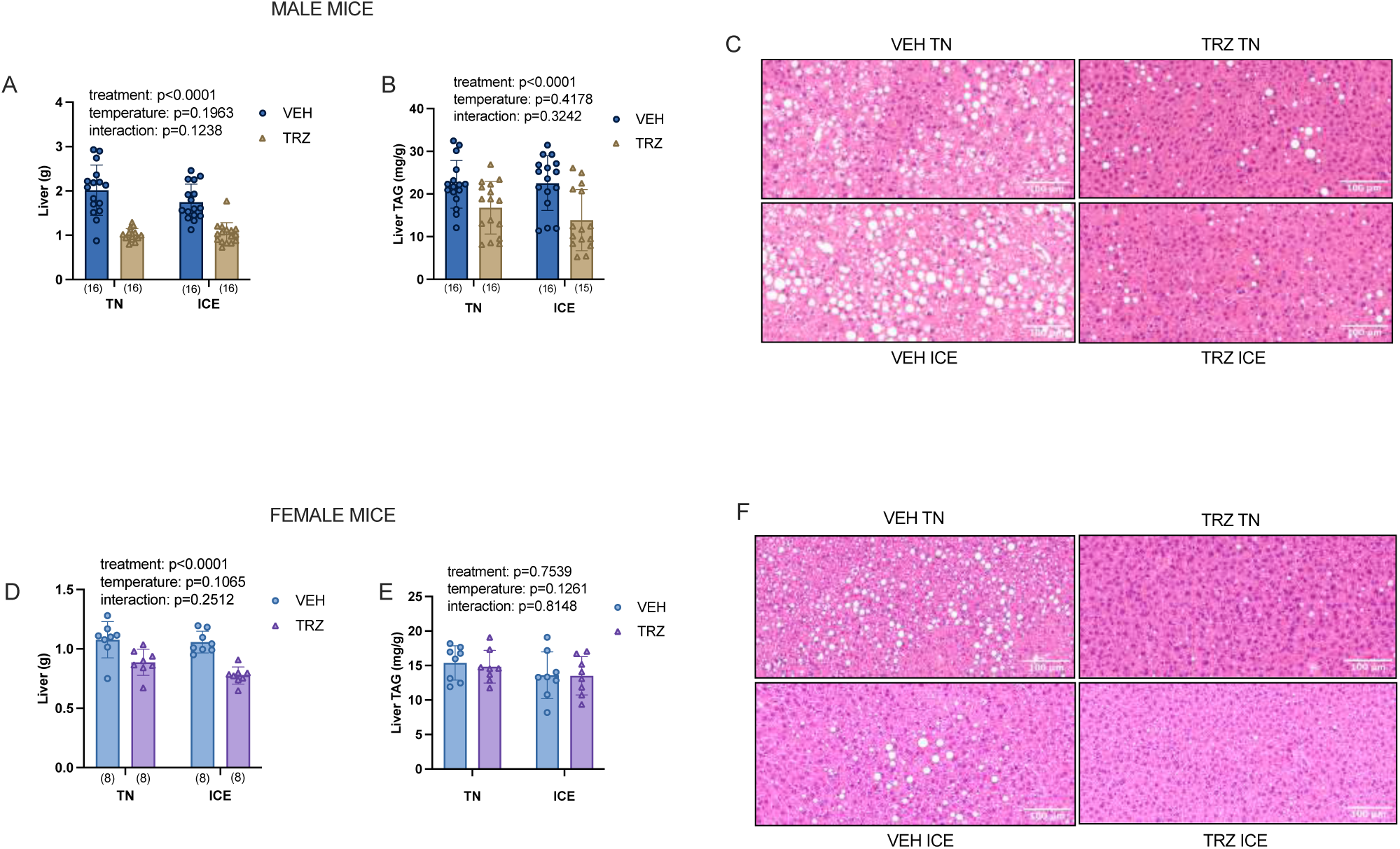
Tirzepatide reduces liver mass and lowers liver triglycerides levels in male mice. Tirzepatide reduces liver mass (A) and liver triglycerides (B) in male mice. In females, tirzepatide lowers liver mass (E), while liver triglycerides were not affected (E). Representative HCE staining images are shown for liver in male (C) and female (F) mice. Statistical analysis was performed using two-way ANOVA (treatment x temperature), with main and interaction effects shown in the inset. Data are presented as mean ± SD, with individual data points shown. Sample sizes per group are indicated in parentheses below each bar.

### ICE and tirzepatide improve glucose homeostasis in DIO mice

Our group has previously shown that ICE improves glucose homeostasis ^24^. In the current study, both tirzepatide and ICE improved glucose tolerance in male mice (Figures 6A and 6B) (treatment: p < 0.0001; temperature: p < 0.0001; interaction: p = 0.1946). The ITT revealed a treatment x temperature interaction in male mice (Figures 6C and 6D) (treatment: p < 0.0001; temperature: p = 0.0262; interaction: p = 0.0465). Post-hoc analyses showed VEH ICE mice had greater AUC than TRZ ICE (p < 0.0001) and TRZ TN (p < 0.0001), and VEH TN mice displayed greater AUC than VEH ICE (p = 0.0163), TRZ TN (p < 0.0001), and TRZ ICE (p < 0.0001). When ITT data were baseline-normalized (0 min = 100), only a main effect of treatment was observed in male mice (figure 6E) (treatment: p = 0.0397; temperature: p = 0.2404; interaction: p = 0.8699).

**Figure 6:**
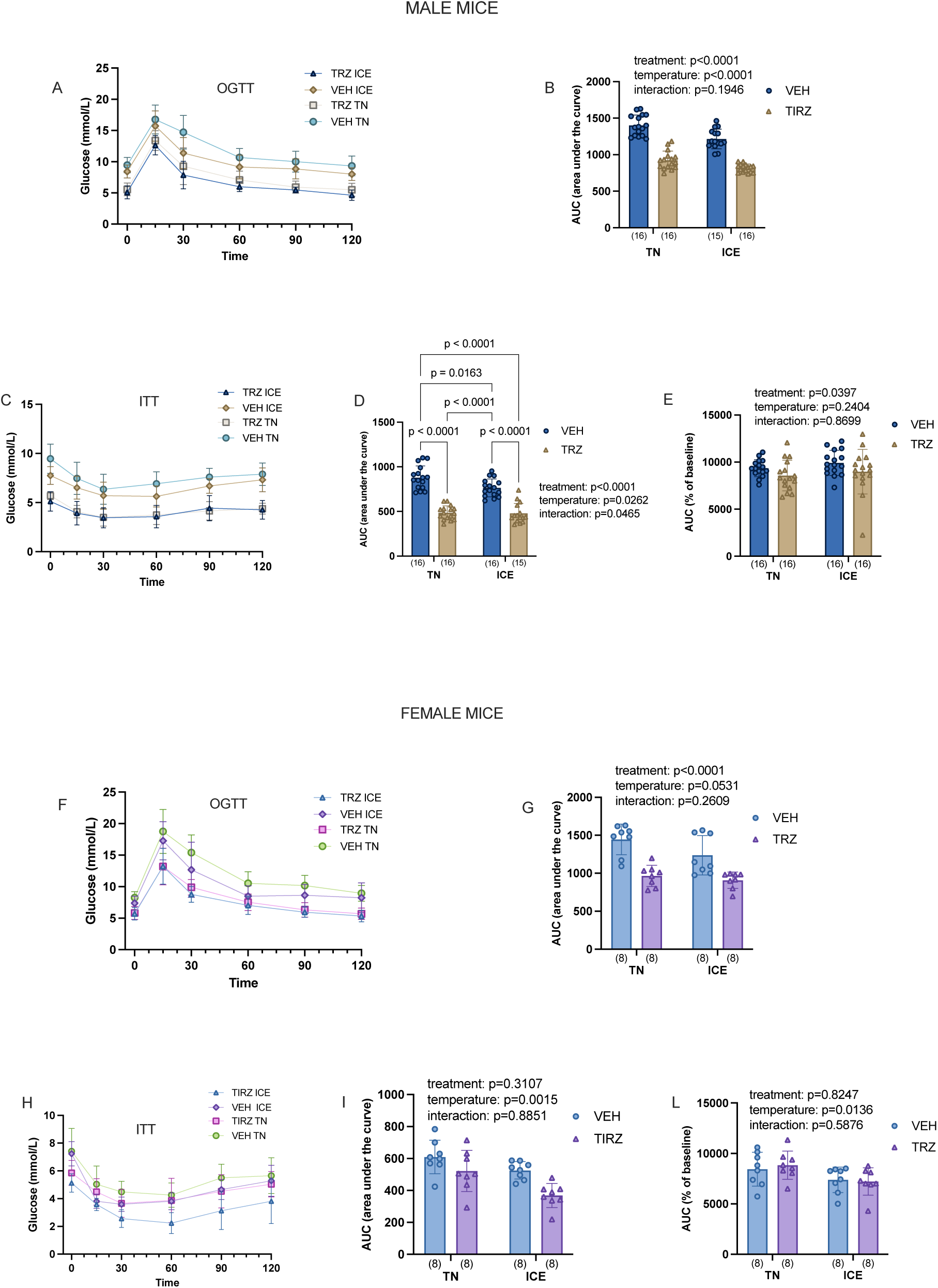
ICE and tirzepatide improve glucose homeostasis in DIO mice. ICE and tirzepatide improve glucose tolerance (A, B) and insulin responsiveness (C, D, E) in male mice. In female mice, tirzepatide improves glucose tolerance (F, G), with a trend toward improvement with ICE, while insulin tolerance is improved with ICE (H, I, L). Panel (E) and (L) show baseline-normalized AUC for the ITT. Statistical analysis was performed using two-way ANOVA (treatment x temperature), with main and interaction effects shown in the inset. Data are presented as mean ± SD, with individual data points shown. Sample sizes per group are indicated in parentheses below each bar.

In female mice, tirzepatide improved glucose tolerance with a trend toward improvements wiht ICE (Figures 6F and 6G) (treatment: p < 0.0001; temperature: p = 0.0531; interaction: p = 0.2609). The ITT showed a main effect of temperature on insulin responsiveness (Figures 6H and 6I) (treatment: p = 0.3107; temperature: p = 0.0015; interaction: p = 0.8851), and a main effect of temperature was also observed when ITT data were baseline-normalized (figure 6L) (treatment: p = 0.8247; temperature: p = 0.0136; interaction: p = 0.5876). Taken together, tirzepatide and ICE improved glucose tolerance in male mice, while in females tirzepatide improved glucose tolerance and ICE showed a tendency to improve glucose handling. For insulin responsiveness, males showed treatment-dependent improvements, whereas in females there was a main effect of ICE.

### Tirzepatide lowers serum cholesterol and fasting insulin levels in DIO male mice

In male mice (Table 1) tirzepatide reduced serum cholesterol (treatment: p = 0.0023; temperature: p = 0.6047; interaction: p = 0.9075) and fasting insulin levels (treatment: p = 0.0038; temperature: p = 0.3212; interaction: p = 0.3932). No differences were observed in serum NEFA (treatment: p = 0.3455; temperature: p = 0.3287; interaction: p = 0.5382), BHB (treatment: p = 0.3193; temperature: p = 0.3213; interaction: p = 0.1503) or triglyceride concentrations (treatment: p = 0.4012; temperature: p = 0.0676; interaction: p = 0.9075). In female mice (Table 2), tirzepatide lowered serum triglyceride levels (treatment: p = 0.0002; temperature: p = 0.3670; interaction: p = 0.0657), and ICE lowered fasting serum insulin levels (treatment: p = 0.1938; temperature: p = 0.0264; interaction: p = 0.7650), with a trend toward greater serum cholesterol with ICE (treatment: p = 0.6610; temperature: p = 0.0546; interaction: p = 0.6830). A treatment x temperature interaction was observed for serum BHB levels (treatment: p = 0.0048; temperature: p = 0.9622; interaction: p = 0.0467). Specifically, TRZ ICE mice display higher serum BHB levels compared to VEH ICE (p = 0.0057).

**Table 1:** Tirzepatide lowers serum cholesterol and fasting insulin levels in DIO male mice. Serum hormones and metabolites were measured in male mice treated with tirzepatide (TRZ) or vehicle (VEH) and housed either at thermoneutrality (TN) or subjected to Intermittent Cold Exposure (ICE). Group sizes are indicated in parentheses. Data are presented as mean ± SD. Statistical differences were assessed using two-way ANOVA (treatment x temperature), and p-values for main and interaction effects are reported.

|  | TRZ TN | VEH TN | TRZ ICE | VEH ICE | p-value |
| --- | --- | --- | --- | --- | --- |
| Insulin (µg/L) | 4.098±1.918(16) | 4.830±1.382(16) | 3.075±1.839(16) | 4.614±1.469(16) | Treatment: 0.0038<br>Temperature: 0.3212<br>Interaction: 0.3932 |
| NEFA (mmol/L) | 0.3248±0.0985(15) | 0.3769±0.1245(15) | 0.3781±0.1181(15) | 0.3891±0.1669(14) | Treatment: 0.3455<br>Temperature: 0.3287<br>Interaction: 0.5382 |
| BHB (mmol/L) | 0.2781±0.1128(16) | 0.2917±0.1060(16) | 0.2918±0.1419(16) | 0.2180±0.1050(14) | Treatment: 0.3193<br>Temperature: 0.3213<br>Interaction: 0.1503 |
| Cholesterol (mM) | 5.490±2.345(15) | 8.720±3.109(15) | 6.154±2.724(16) | 7.334±2.571(15) | Treatment: 0.0023<br>Temperature: 0.6047<br>Interaction: 0.1442 |
| Triglycerides (mg/dL) | 66.63±11.53(15) | 70.24±8.696(16) | 74.06±18.71(16) | 76.79±17.45(14) | Treatment: 0.4012<br>Temperature: 0.0676<br>Interaction: 0.9075 |

**Table 2:** Tirzepatide and ICE differentially affect circulating triglycerides, BHB, and insulin levels in female DIO mice. Serum hormones and metabolites were measured in female mice treated with tirzepatide (TRZ) or vehicle (VEH) and housed either at thermoneutrality (TN) or subjected to Intermittent Cold Exposure (ICE). Group sizes are indicated in parentheses. Tukey’s post-hoc test was applied when a significant treatment x temperature interaction occurred. Letters indicate significant post-hoc comparison. Data are presented as mean ± SD. Statistical differences were assessed using two-way ANOVA (treatment x temperature), and p-values for the main and interaction effects are reported.

|  | TRZ TN | VEH TN | TRZ ICE | VEH ICE | p-value |
| --- | --- | --- | --- | --- | --- |
| **Insulin**<br>($\mu\text{g/L}$) | 1.219±0.9392(8) | 1.422±0.5062(8) | 0.6970±0.28918) | 1.019±0.1548(8) | Treatment: 0.1938<br>Temperature: 0.0264<br>Interaction: 0.7650 |
| **NEFA**<br>(mmol/L) | 0.43778±0.1378(8) | 0.3833±0.1036(7) | 0.3333±0.1195(7) | 0.4353±0.1531(8) | Treatment: 0.3455<br>Temperature: 0.3287<br>Interaction: 0.5382 |
| **BHB**<br>(mmol/L) | 0.3166±0.1692(8) | 0.2751±0.0616(16) | <sup>a</sup> 0.4022±0.1513(8) | <sup>a</sup> 0.1855±0.0371(8) | Treatment: 0.0048<br>Temperature: 0.9622<br>Interaction: 0.0467 |
| **Cholesterol**<br>(mM) | 2.180±0.7676(8) | 2.168±0.5280(8) | 3.074±1.527(8) | 2.758±1.085(8) | Treatment: 0.6610<br>Temperature: 0.0546<br>Interaction: 0.6830 |
| **Triglycerides**<br>(mg/dL) | 16.41±1.111(7) | 20.08±3.815(7) | 12.22±3.774(7) | 21.56±5.506(8) | Treatment: 0.0002<br>Temperature: 0.3670<br>Interaction: 0.0657 |

### Tirzepatide and ICE differentially affect circulating triglycerides, BHB, and insulin levels in female DIO mice

In female mice (Table 2), tirzepatide lowered serum triglyceride levels (treatment: p = 0.0002; temperature: p = 0.3670; interaction: p = 0.0657), and ICE lowered fasting serum insulin levels (treatment: p = 0.1938; temperature: p = 0.0264; interaction: p = 0.7650), with a trend toward greater serum cholesterol with ICE (treatment: p = 0.6610; temperature: p = 0.0546; interaction: p = 0.6830). A treatment x temperature interaction was observed for serum BHB levels (treatment: p = 0.0048; temperature: p = 0.9622; interaction: p = 0.0467). Specifically, TRZ ICE mice display higher serum BHB levels compared to VEH ICE (p = 0.0057).

### Tirzepatide does not enhance energy expenditure in mice housed at thermoneutrality

Tirzepatide has been shown to impact energy expenditure and substrate utilizations ^35^. In our study, we found that in mice housed at thermoneutrality, tirzepatide did not affect total energy expenditure, but there was a main effect of body mass (figure 7B and 7C) (mass: p < 0.0001; group: p = 0.8099). No differences were observed during the dark phase (figure 7C) (mass: p < 0.0001; group: p = 0.4307) or the light phase (figure 7C) (mass: p < 0.0001; group: p = 0.8150). At thermoneutrality, average RER over 24 hours was decreased with tirzepatide (figure 7D) (treatment: p < 0.0001; temperature: p = 0.0620; interaction: p = 0.1428), an effect seen in both the dark phase (figure 7D) (treatment: p = 0.0001; temperature: p = 0.4064; interaction: p = 0.1876) and the light phase (figure 7D) as well (treatment: p = 0.0005; temperature: p = 0.9244; interaction: p = 0.0729). There were no differences in total locomotion (figure 7E) (treatment: p = 0.5178; temperature: p = 0.8595; interaction: p = 0.6571). Fat oxidation at thermoneutrality was not affected by tirzepatide (figure 7F) (treatment: p = 0.5640; temperature: p = 0.3785; interaction: p = 0.2954), nor was carbohydrate oxidation (figure 7G) (treatment: p = 0.1637; temperature: p = 0.5981; interaction: p = 0.2073).

**Figure 7:**
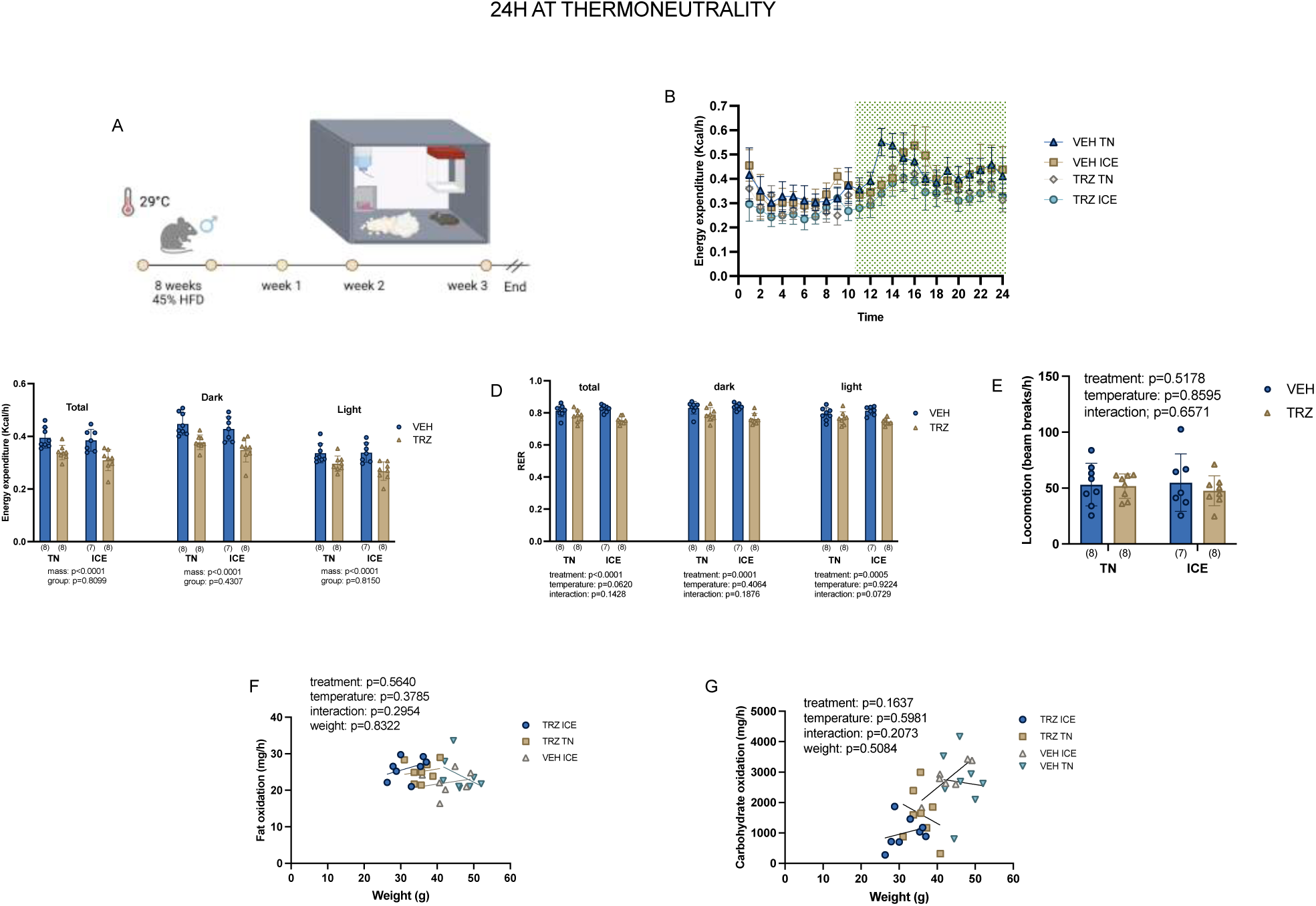
Tirzepatide does not enhance energy expenditure in mice housed at thermoneutrality. Tirzepatide and ICE had no effect on energy expenditure (B, C) locomotion (E), fat oxidation (F), or carbohydrate oxidation (G), but lowered RER (D) in male mice housed at thermoneutrality. Statistical analyses were performed using a linear regression model including treatment, temperature, their interaction, and body weight as a covariate. Main and interaction effects are shown in the inset. Data are presented as mean ± SD, with individual data points shown. Sample sizes per group are indicated in parentheses below each bar.

### ICE acutely increases energy expenditure

We assessed energy expenditure and substrate utilization during a 1 h cold challenge. Energy expenditure increased with cold exposure (figure 8A and 8B) (treatment: p = 0.9364; temperature: p < 0.0001; interaction: p = 0.0408). A treatment x temperature interaction was observed (figure 8B), with TRZ ICE mice displaying higher energy expenditure compared to TRZ TN (p < 0.0001), and VEH ICE mice showing higher energy expenditure compared to VEH TN (p < 0.0001). Moreover, fat oxidation increased with ICE (figure 8C and 8D) (treatment: p = 0.8321; temperature: p < 0.0001; interaction: p = 0.5350), whereas carbohydrate oxidation did not change during the 1 h cold exposure (figure 8E and 8F) (treatment: p = 0.7062; temperature: p = 0.8208; interaction: p = 0.9633). Overall, tirzepatide did not impact the ability of ICE to increase energy expenditure and fat oxidation.

**Figure 8:**
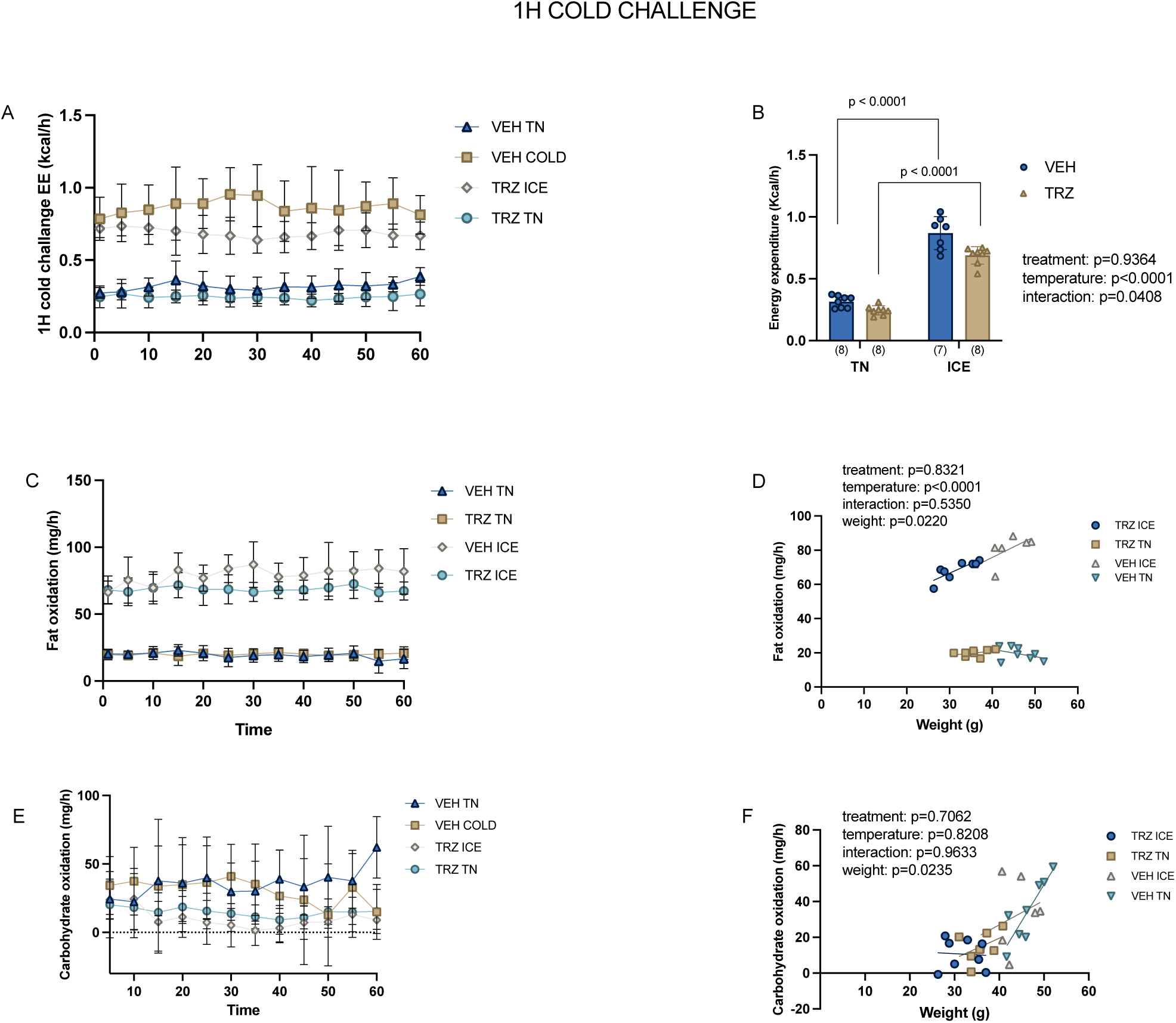
ICE acutely increases energy expenditure. ICE increased energy expenditure (A, B) and fat (C, D) but not carbohydrate oxidation (E,F). Statistical analyses were performed using a linear regression model including treatment, temperature, their interaction, and body weight as a covariate. Main and interaction effects are shown in the inset. Data are presented as mean ± SD, with individual data points shown. Sample sizes per group are indicated in parentheses below each bar.

## Discussion

In the present study, we investigated the individual and combined effects of tirzepatide and ICE on body composition, energy expenditure, and glucose homeostasis in DIO mice housed at thermoneutrality. Our findings demonstrate that tirzepatide effectively reduced body weight and adiposity and improved glucose homeostasis while ICE treatment, independent of changes in body weight, improved glucose homeostasis. Tirzepatide, was more effective at treating obesity and dysregulated glucose metabolism than ICE and there was no additional benefit of combining the two interventions.

While effective at causing weight loss, there has been concern raised about the loss of lean tissue with GLP1RAs^36^. In the current study we found that tirzepatide reduced lean and skeletal muscle mass in male but not female mice. These findings are similar to a recent pre-print demonstrating that female *ob/ob* mice are protected against semaglutide induced muscle loss^37^ and are consistent with evidence that males lose proportionally more weight and lean mass during caloric restriction compared to females^38^. As cold exposure can induce shivering-related muscle contractions^39^, it is somewhat surprising that ICE did not attenuate the reductions in lean mass observed with tirzepatide. It is possible that a more robust induction of muscle contractions, whether through shivering or other interventions, may be required to preserve lean mass.

Previous studies have examined the impact of GLP1-RAs on energy expenditure and metabolic adaptations in rodents and humans^35^. In our study, tirzepatide did not alter total energy expenditure, fat oxidation, or carbohydrate oxidation at thermoneutrality, suggesting that its metabolic benefits are primarily driven by reduced energy intake rather than changes in energy expenditure. Indeed, it has been shown^35^ that tirzepatide does not impact metabolic adaptation in people with obesity but increases fat oxidation in both humans and mice. In line with this, we observed a reduction in RER in tirzepatide-treated mice housed at thermoneutrality, indicating a shift toward greater lipid utilization.

In contrast to tirzepatide, energy expenditure and fat oxidation were markedly increased during a 1 h cold challenge and this effect was not impacted by tirzepatide treatment. Interestingly, despite causing large increases in energy expenditure, we did not observe ICE-induced reductions in body weight.

In previous work from our group we found that 4 weeks of ICE treatment caused subtle increases in body weight that were driven by increases in food intake in singly housed mice. In the present investigation we did not detect differences in food intake, though it should be noted that mice in this study were group housed and we might lack the sensitivity to detect small changes in food intake. Regardless of these differences, we demonstrate, as before ^24^ an uncoupling between, weight loss and improvements in glucose homeostasis with ICE.

Improvements in glucose tolerance were more robust with tirzepatide compared to ICE in both sexes and there didn’t seem to be any additional benefit of combining both interventions. Conversely, insulin tolerance was improved with tirzepatide in male mice and with ICE in female mice. While the underlying mechanisms explaining these sex differences are not known, sex differences in glucose metabolism have been well documented in C57BL/6J mice^40^, with females exhibiting greater glucose tolerance and insulin responsiveness compared to males^41^. This aligns with the well known influence of sex hormones on glucose metabolism^40, 42^, as well as previously reported sex differences in baseline glucose homeostasis and pancreatic β-cell adaptation^41, 42^

To our knowledge, this is the first study to examine the combined effects of tirzepatide and intermittent cold exposure on body composition, energy metabolism, and glucose homeostasis in both male and female DIO mice housed at thermoneutrality. Housing mice at themroneutrality, more closely reflects the human thermal environment and removes the confounding influence of chronic cold-induced thermogenesis present under standard housing conditions^43^. The observation that ICE improves glucose homeostasis independently of changes in body weight, and that this effect persists despite tirzepatide-induced weight loss, might indicate that cold exposure engages distinct, weight-independent mechanisms of glucose regulation. Further investigation into the role of sex hormones in modulating the interaction between thermoregulatory and incretin signaling will be essential to understand the observed sex-specific responses and to inform more personalized metabolic interventions.

Overall, our findings suggest that targeting energy intake and energy expenditure may represent complementary, but not necessarily additive approaches to improving metabolic health. Tirzepatide primarily improves metabolic outcomes through reductions in food intake and body weight, whereas ICE enhances glucose homeostasis and substrate utilization independently of changes in body weight and adiposity. The absence of additive effects on energy expenditure and related metabolic readouts indicates that pharmacological and environmental interventions may act through distinct physiological pathways.

## Authors contribution

AB and DCW conceived and designed the research. AB, HA, and CG performed daily injections and ICE. AB, MA, and BW analyzed HCE data. AB, HA, CG, KE, BB, and ABM performed OGTT and ITT. AB, HA, MA, KE, and BB performed tissue collection. AB, HA, and SJ conducted metabolic cage experiments. AB and DCW analyzed the data and interpreted the results. AB and DCW prepared the figures, wrote and revised the manuscript. All authors approved the final version of the manuscript.

## References

1. Evans, M. et al. Obesity-related complications, healthcare resource use and weight loss strategies in six European countries: the RESOURCE survey. Int J Obes (Lond) 47, 750–757 (2023).

2. Meneguetti, B.T. et al. Neuropeptide receptors as potential pharmacological targets for obesity. Pharmacol Ther 196, 59–78 (2019).

3. Bluher, M. Obesity: global epidemiology and pathogenesis. Nat Rev Endocrinol 15, 288–298 (2019).

4. Jensterle, M., Rizzo, M., Haluzik, M. & Janez, A. Efficacy of GLP-1 RA Approved for Weight Management in Patients With or Without Diabetes: A Narrative Review. Adv Ther 39, 2452–2467 (2022).

5. D’Alessio, D. Is GLP-1 a hormone: Whether and When? J Diabetes Investig 7 Suppl 1, 50–55 (2016).

6. Huang, Z. et al. Glucose-sensing glucagon-like peptide-1 receptor neurons in the dorsomedial hypothalamus regulate glucose metabolism. Sci Adv 8, eabn5345 (2022).

7. Dong, Y. et al. Time and metabolic state-dependent effects of GLP-1R agonists on NPY/AgRP and POMC neuronal activity in vivo. Mol Metab 54, 101352 (2021).

8. Gallwitz, B. et al. GLP-1-analogues resistant to degradation by dipeptidyl-peptidase IV in vitro. Regul Pept 86, 103–111 (2000).

9. Drucker, D.J. Enhancing incretin action for the treatment of type 2 diabetes. Diabetes Care 26, 2929–2940 (2003).

10. Shah, M. & Vella, A. Effects of GLP-1 on appetite and weight. Rev Endocr Metab Disord 15, 181–187 (2014).

11. Ussher, J.R. & Drucker, D.J. Glucagon-like peptide 1 receptor agonists: cardiovascular benefits and mechanisms of action. Nat Rev Cardiol 20, 463–474 (2023).

12. Frias, J.P. Tirzepatide: a glucose-dependent insulinotropic polypeptide (GIP) and glucagon-like peptide-1 (GLP-1) dual agonist in development for the treatment of type 2 diabetes. Expert Rev Endocrinol Metab 15, 379–394 (2020).

13. Willard, F.S., et al. Tirzepatide is an imbalanced and biased dual GIP and GLP-1 receptor agonist. JCI Insight 5 (2020).

14. von Degenfeld, G., Wehrman, T.S., Hammer, M.M. & Blau, H.M. A universal technology for monitoring G-protein-coupled receptor activation in vitro and noninvasively in live animals. FASEB J 21, 3819–3826 (2007).

15. Nauck, M.A. & D’Alessio, D.A. Tirzepatide, a dual GIP/GLP-1 receptor co-agonist for the treatment of type 2 diabetes with unmatched effectiveness regrading glycaemic control and body weight reduction. Cardiovasc Diabetol 21, 169 (2022).

16. Regmi, A. et al. Tirzepatide modulates the regulation of adipocyte nutrient metabolism through long-acting activation of the GIP receptor. Cell Metab 36, 1534–1549 e1537 (2024).

17. Samms, R.J. et al. GIPR agonism mediates weight-independent insulin sensitization by tirzepatide in obese mice. J Clin Invest 131 (2021).

18. Ivanova, Y.M. & Blondin, D.P. Examining the benefits of cold exposure as a therapeutic strategy for obesity and type 2 diabetes. J Appl Physiol (1985) 130, 1448–1459 (2021).

19. Lee, P. et al. Temperature-acclimated brown adipose tissue modulates insulin sensitivity in humans. Diabetes 63, 3686–3698 (2014).

20. Bing, C. et al. Hyperphagia in cold-exposed rats is accompanied by decreased plasma leptin but unchanged hypothalamic NPY. Am J Physiol 274, R62–68 (1998).

21. Shefer, V.I. & Talan, M.I. Change in heat loss as a part of adaptation to repeated cold exposures in adult and aged male C57BL/6J mice. Exp Gerontol 32, 325–332 (1997).

22. Talan, M.I., Engel, B.T. & Whitaker, J.R. A longitudinal study of tolerance to cold stress among C57BL/6J mice. J Gerontol 40, 8–14 (1985).

23. Ravussin, Y., Xiao, C., Gavrilova, O. & Reitman, M.L. Effect of intermittent cold exposure on brown fat activation, obesity, and energy homeostasis in mice. PLoS ONE 9, e85876 (2014).

24. McKie, G.L. et al. Intermittent cold exposure improves glucose homeostasis despite exacerbating diet-induced obesity in mice housed at thermoneutrality. J Physiol 600, 829–845 (2022).

25. Samms, R.J. et al. Tirzepatide induces a thermogenic-like amino acid signature in brown adipose tissue. Mol Metab 64, 101550 (2022).

26. Coskun, T. et al. LY3298176, a novel dual GIP and GLP-1 receptor agonist for the treatment of type 2 diabetes mellitus: From discovery to clinical proof of concept. Mol Metab 18, 3–14 (2018).

27. Bellucci, A. et al. Topical application of the cold-mimetic l-menthol decreases wheel running without affecting the beneficial effects of voluntary exercise in mice. Exp Physiol (2025).

28. Nagy, C. & Einwallner, E. Study of In Vivo Glucose Metabolism in High-fat Diet-fed Mice Using Oral Glucose Tolerance Test (OGTT) and Insulin Tolerance Test (ITT). J Vis Exp (2018).

29. Weir, J.B. New methods for calculating metabolic rate with special reference to protein metabolism. J Physiol 109, 1–9 (1949).

30. Girousse, A. et al. Surplus fat rapidly increases fat oxidation and insulin resistance in lipodystrophic mice. Mol Metab 13, 24–29 (2018).

31. Mina, A.I. et al. CalR: A Web-Based Analysis Tool for Indirect Calorimetry Experiments. Cell Metab 28, 656–666 e651 (2018).

32. Jeromson, S. et al. Semaglutide impacts skeletal muscle to a similar extent as caloric restriction in mice with diet-induced obesity. J Physiol (2025).

33. Look, M. et al. Body composition changes during weight reduction with tirzepatide in the SURMOUNT-1 study of adults with obesity or overweight. Diabetes Obes Metab 27, 2720–2729 (2025).

34. Li, Y., Sun, W., Liu, H. & Ruan, X.Z. Tirzepatide, a dual GIP/GLP-1 receptor agonist, alleviates metabolic dysfunction-associated steatotic liver disease by reducing the expression of CD36 and OBP2A. Genes Dis 12, 101761 (2025).

35. Ravussin, E. et al. Tirzepatide did not impact metabolic adaptation in people with obesity, but increased fat oxidation. Cell Metab 37, 1060–1074 e1064 (2025).

36. Neeland, I.J., Linge, J. & Birkenfeld, A.L. Changes in lean body mass with glucagon-like peptide-1-based therapies and mitigation strategies. Diabetes Obes Metab 26 Suppl 4, 16–27 (2024).

37. Rout, S., et al. Females are protected from semaglutide-induced muscle loss in ob/ob mice. bioRxiv (2026).

38. de Souza, G.O., Wasinski, F. & Donato, J., Jr. Characterization of the metabolic differences between male and female C57BL/6 mice. Life Sci 301, 120636 (2022).

39. Blondin, D.P. & Haman, F. Shivering and nonshivering thermogenesis in skeletal muscles. Handb Clin Neurol 156, 153–173 (2018).

40. Gao, A. et al. Sexual dimorphism in glucose metabolism is shaped by androgen-driven gut microbiome. Nat Commun 12, 7080 (2021).

41. Mastrolia, V. et al. Loss of alpha(2)delta-1 Calcium Channel Subunit Function Increases the Susceptibility for Diabetes. Diabetes 66, 897–907 (2017).

42. Jacobo-Piqueras, N., Theiner, T., Geisler, S.M. & Tuluc, P. Molecular mechanism responsible for sex differences in electrical activity of mouse pancreatic beta cells. JCI Insight 9 (2024).

43. Bellucci, A., Baranowski, B.J. & Wright, D.C. The effects of housing temperature on mouse physiology and implications for disease modeling. Lab Anim (NY) 55, 108–116 (2026).

